# Specific Ethogram of the Mexican four-eyed octopus: *Octopus maya*

**DOI:** 10.1101/2022.10.10.511610

**Authors:** D.A. González-Navarrete, F. Vergara-Ovalle, P. García-Andaluz, F. Ayala-Guerrero, C. Rosas, P. Vázquez-León, D.B. Paz-Trejo, H. Sánchez-Castillo

## Abstract

Historically, behavior studies have focused mainly on animals of two phyla in particular: Craniata and Arthropoda, however, behavioral research on alternative phyla like mollusks has been increasing because of the potential for research that these models present. When we talk about mollusk behavior, cephalopods are the first group that stands out, however, research on Mexico’s endemic species like *Octopus maya*, is still lacking. This octopus could help to reach a standardized model in neuroscience, because adapts well to laboratory conditions and has been successfully cultured through several generations. These characteristics provide a great advantage as a research model since they could reduce the number of variables that affect behavioral studies, something hard to control with a captured-wild octopus. Hence, in order to work properly with species like this, in research environments, it’s fundamental to know first the behaviors that this species can perform there. Therefore, the objective of this study was to elaborate an ethogram that describes the behavioral repertoire that *O. maya* displays in laboratory conditions. Thirteen individuals of *O. maya* (6-20g) were used and maintained in tanks with a closed circulation seawater system and illuminated with a red light of 30 lx in a 12:12 LD cycle. Nine of these individuals were used for an *ad libitum* sampling of behaviors to name, define, categorize and operationally describe them. The last four individuals were used to establish day/night activity patterns, (length and frequency of behaviors throughout the day). The obtained results showed that *O. maya* has a wide behavioral repertoire with at least twenty-three behaviors, which were included in six different behavioral categories. The obtained results showed that *O. maya* has a wide behavioral repertoire with at least twenty-three behaviors, which were organized in six different behavioral categories. Also, this species showed differentiated activity patterns between day and night, with peaks of activity, distribution, and frequencies of activity behaviors mainly during the light hours. These results showed that *O. maya* has behaviors comparable with other octopus species and support the proposal for its use as a viable research model. Knowing the behavioral repertoire of *O. maya* allows for better control in future behavioral studies using this model, provides the main tools to evaluate the organism’s health and status, and supports its use for research in neuroscience and cognition.

## INTRODUCTION

Behavior is a phenomenon that has been studied from different fields and approaches including Neuroscience, Ethology, and Psychology. Ethogram is one of the most important tools for the study of behavior, it consists of a list of specific behaviors that a species can perform in a specific environment and works as an observational instrument to collect information (Lehner, 1996; Riba, 1988). In general, an ethogram aims to give a complete picture of the behavior from an evolutionary perspective, answering the questions of what, who, how, where, and why of a given behavior, producing a replicable starting point for behavioral research.

Historically, behavior studies have focused mainly on animals of two phyla in particular: Craniata (where we mainly find studies in mammals) and Arthropoda (mainly insect studies), however, the study of behavior on alternative phyla has been increasing (Schnell & Clayton, 2021). For example, behavioral studies with mollusks, particularly cephalopods, stand out for their potential for research (Mather, 2008). The class Cephalopoda (nautilus, squids, cuttlefish, and octopus) is considered the most complex one in the phylum Mollusca (Fiorito et al., 2014), and one of the most studied cephalopods in neuroscience is octopuses (Borrelli & Fiorito, 2008). There are nearly 300 recognized species of octopus and all of them present a similar anatomic structure: the mantle on the top of its body where they have enclosed the internal organs and eight arms on the bottom with 250 suckers each, that are connected to the head, composed by the eyes, the mouth and the cerebral ganglia (O’Brien et al., 2018). Furthermore, octopuses possess a nervous system with more than 500 million neurons, of which approximately 200 million are centralized in the cerebral ganglia. These ganglia in the head are composed of three main parts: supra-esophageal mass, sub-esophageal mass, and two optic lobes (Borrelli & Fiorito, 2008; Di Cosmo et al., 2018; O’Brien et al., 2018); those structures receive, integrate and process sensorial cues to perform an appropriated behavior. This particular arrangement provides octopuses with more complex cognitive skills (attentional, learning and memory, etc.) than other invertebrates (Grasso & Basil, 2009; O’Brien et al., 2018). Their complex nervous system, alongside their efficient physiology and motor control, form the basis for complex behavior and allow them to perform a wide behavioral repertoire (Hochner, 2008; Mather, 2008), being a good candidate for a wide range of neuroscience studies (behavioral, ethological, molecular, cellular, etc).

The regular behaviors that some species of octopus perform have been described in a few specific publications (Borrelli et al., 2006; Hanlon & Messenger, 1996; Mather, 2010; Mather & Scheel, 2014; Wells, 1978), and to our knowledge, there is a lack description of the entire behavioral repertoire of the octopus. On this subject we found two main studies: the first one describes a specific ethogram for *Abdopus aculeatus* (Huffard, 2007), and the second one describes some general behaviors of seventeen members of the Octopodidae family (Mather & Alupay, 2016). Other articles have described individual behaviors of different octopuses (*O. vulgaris*, *Eledone cirrhosa*, and *O. bimaculoides)*, among which are the description of components, postures, and actions in the use of arms (Mather, 1998), feeding and foraging behaviors (Mather, 1991a), patterns of coloration and movement of the whole body after exposure to an aversive stimulus (Packard & Sanders, 1971), den occupancy, maintenance and blocking behaviors (Mather, 1994), attack and withdrawal behaviors related to den use (Cigliano, 1993), inactivity, activity and alerting behaviors (Cobb et al., 1995) and sleeping, resting, hunting, feeding, den maintenance, and grooming (Mather, 1988).

Nevertheless, despite the fact that isolated descriptions of some behaviors provide us with great information about octopuses, the evidence can’t generalize too easily from one species to another because may there slight differences between them due to they move and adapt in a wide variety of marine habitats with different characteristics, which could modulate some specific behaviors related to the habitat but not a general behavior (Mather & Scheel, 2014; O’Brien et al., 2018). All this clarifies the need for a correct standardization of behavioral evaluation, in order to achieve this, it’s necessary for the initial study of the complete behavioral repertoire of the species investigated to get a particular point of reference about the organism.

There are important reports on the behavior of octopuses, but it is not yet possible to perform a standardization for the evaluation of behavior because the information available has been collected in different conditions and different species of octopuses, many of them lacking controlled experimental observations on the behavioral repertoire (Borrelli & Fiorito, 2008). Those circumstances around data could be the source of the variability found in behavioral studies on octopuses (Borrelli et al., 2020). Besides, the limitation implicated in the use of captured-wild octopus brings another set of complications as the lack of control of variables such as the life story of the subjects or previous exposure to aversive stimuli. If those variables are not controlled or known, they could affect behavioral performance in unforeseen ways and produce different responses to the same stimulus. In the aim to diminish the impact of these variables, the use of octopus species that are adapted to laboratory conditions and that can be kept in captivity from hatching, become a great candidate for the study of behavior in these organisms, providing an optimal reference point for neuroscience research.

In this sense, *Octopus maya*, also known as the Mexican four-eyed octopus (Fig. 1a), is an endemic species of the Yucatán peninsula present in shallow waters of the states of Campeche, Yucatán, and Quintana Roo. This octopus is characterized by dark brown skin, either in a lighter tone or in a reddish tone in the presence of aversive stimuli (Voss & Solis-Ramirez, 1966); also presents two black spots, one under each eye, between the bases of the second and third arm. One main characteristic of the use of this species of octopus in neuroscience research it’s because has adapted very well to laboratory conditions, successfully achieving breeding in captivity through several generations (Hanlon & Forsythe, 1985; Rosas et al., 2007; Van Heukelem, 1976). Currently, captive breeding in México is being developed by Universidad Nacional Autónoma de México (UNAM) at the Unidad Multidisciplinaria de Docencia e Investigación in Sisal, Yucatán since 2004 (Rosas et al., 2014), providing this species with a great advantage as a research model for octopus’s behavior, reducing the number of variables that could affect the behavioral studies.

**Figure 2.**
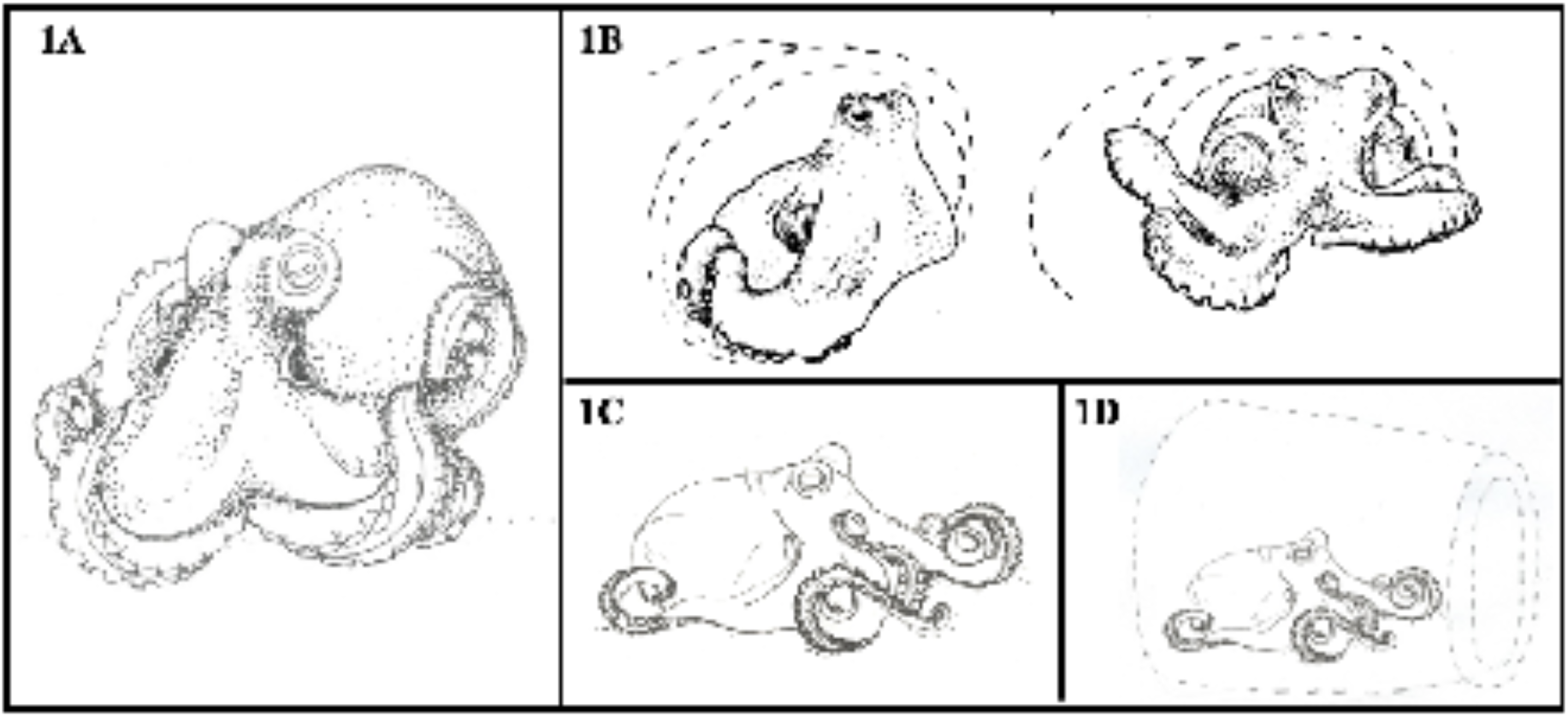
Rest behaviors of *O. maya*. A) Drawing of *Octopus maya*. B) Rest within the den: No activity inside the den. C) Rest outside the den: No activity outside the den. D) Total rest: Completely inside of den with no visible activity

Thus, working with a species of octopus with these characteristics will allow us a proper interpretation of the results and changes observed, ensuring that these derive from the applied experimental conditions and not only due to the organism’s life history. In order to be able to work properly with a species like this in laboratory conditions, it is necessary to know in detail its full behavioral repertoire in said conditions. Taking all this into consideration, the aim of this study was to construct a specific ethogram of the behaviors displayed by juvenile organisms of the species *O. maya* raised and maintained in laboratory conditions, through the techniques of observation and description provided by ethology. This ethogram could establish a behavioral benchmark for future neuroscience research through a better understanding of octopus behavior.

## MATERIALS AND METHODS

### Subjects and housing

A total of 13 subjects of O. maya of two months old (6-20g) were observed. All octopuses were bred in captivity and obtained from the Laboratory of Applied Ecophysiology in the Multidisciplinar Unit of Teaching and Research located in Sisal Yucatán, México. The specimens were sent and transported to the Psychology School UNAM in México City. The octopuses were housed in two large tanks of 120cm x 40cm x 40cm (length, width, and height respectively) with a capacity of 2 specimens each. The tanks contained natural sand or gravel substrate, shells of various sizes, and a clay vase and were illuminated by a 30-lx red light in a 12-12 light-dark cycle, in accordance with the species’ natural environment ecology. Weekly water changes and full monthly cleaning procedures of tanks were performed. The sex of the organisms was not determined since at this age they have not reached sexual maturity yet, and the presence of sexual dimorphism is not yet present.

Artificial salt water (Instant Ocean) was prepared and kept in constant circulation through a closed circulation system, with an attached external filter with skimmers, UV filters, and sponges with denitrifying bacteria. Conditions were maintained at standard salinity levels of 3.5%, pH of 7.8-8.3, temperature of 25 °C ± 2 °C, and low nitrite and ammonium levels (between 0.0 −0.1). All these parameters in the tanks were evaluated every third day during the first 2 weeks of acclimatization and weekly during the rest of the housing time. The specimens were fed with frozen pieces of shrimp or squid tentacles twice a day (10:00 am and 6:00 pm).

### Acclimatization

A two-week acclimatization period was allowed to avoid stress caused by transportation and handling procedures. We establish a parameter of total acclimatization to the environment the appropriation of some space or object within the tank where the specimen spends most of the time and uses it as a den. The health of the specimens was monitored according to the Guidelines for the Care and Welfare of Cephalopods in Research (Fiorito, et al., 2015).

### Behavioral record

Two types of behavioral recordings were performed. In the first place, nine specimens were used for an *ad libitum* sampling of the behavioral repertoire through short videos (5 minutes-10 minutes) obtained by using a camera handled by the experimenter and a constant following of octopus movements during daylight hours. This recording was made to obtain a catalog and general characterization of all behaviors presented by the specimens.

In the second place, four different specimens were used to determine periods of activity, frequency, and duration of behaviors displayed, through a continuous recording of 24 hours without the presence of researchers (with the exception of feeding periods) and with the red light constantly turned on. For this, a full HD video camera (Sony, Inc, USA) was mounted on a tripod 70 cm away from the tanks and connected to a computer for storage and posterior analysis of videos.

### Behavioral analysis

Observed behaviors from the *ad libitum* sampling were named, categorized, and operationally defined. Behaviors were included into six different behavioral categories (rest, locomotion, feeding, home maintenance, protection, and others) and representative postures of each were illustrated by drawings for a better understanding of taking the postures of each behavior to do that. Postures were considered as a constant body configuration maintained over time, either for a long time as in resting behaviors or for a short time as in activity behaviors, which allows better identification of the behavior that the organism is displaying.

The previously described behaviors were used to analyze the 24 hours continuous videos of four recorded octopuses. The duration of behaviors was grouped in 10-minute intervals for the analysis in order to obtain the periods of activity and non-activity during the 24 hours. Also, the frequency of appearance of each behavior was grouped to know which activity behaviors occur most frequently throughout the day.

### Statistical analysis

Inter-observer reliability of analyzes was obtained using Cohen’s Kappa coefficient for all videos. On the other hand, to determine differences in activity periods, the 24 hours were divided into the hours corresponding to the day (12 hours) and the hours corresponding to the night (12 hours) and a nonparametric Wilcoxon sign classification test was run. On the other hand, to determine differences in activity periods, the 24 hours were divided into the hours corresponding to the day (12 hours) and the hours corresponding to the night (12 hours) and a nonparametric Wilcoxon sign classification test was run. Finally, to determine the most frequent behaviors throughout 24 hours, we run a non-parametric test of Friedman’s ANOVA followed by an uncorrected Dunn’s multiple comparison test of all activity-related behaviors, to identify differences between each of them.

## RESULTS

A value of 0.612 was obtained after applying a Cohen’s kappa coefficient analysis on the videos obtained from the four specimens (1B, 2B, 2A, and 1A), which indicates a good level of agreement.

## DESCRIPTION OF THE BEHAVIORS

The ethogram of *O. maya* in laboratory conditions is composed of twenty-three behaviors divided into six behavioral categories (Table 1). Due to insufficient evidence in the literature on the behavior of *O. maya*, the descriptions were made only in a structural way (except in the category of feeding and protection). The definition of each of them, as well as an illustrative image representing the characteristic posture of the presented behavior.

**Table 1.** Behavioral categories of *O. maya* and their respective behaviors.

| **Behavioral categories of *O. maya*** |  |  |  |  |  |
| --- | --- | --- | --- | --- | --- |
| **Rest** | **Locomotion** | **Feeding** | **Home Maintenance** | **Protective** | **Others** |
| Within den | Crawling | Hunt | Excavation | Isolation | Defecation |
| Outside den | Hovering | Reach | Manipulate | Inking | Grooming |
| Total | Jet Propulsion | Feed |  | Protection | Rearing |
|  | Climbing | Waste disposal |  | Push |  |
|  | Jump on shells |  |  | Flatten |  |
|  |  |  |  | Hand fan |  |

### Rest behaviors

Behaviors related to the non-activity of the specimen, are sometimes accompanied by a rearrangement of the body, both inside and outside the den (Fig. 1).

#### Rest Within the Den

In this observed behavior, the octopus is partially inside the den with its front arm arranged in a mustache-like form, in a way that the mantle and half of the arms are inside while the head and the other half of the arms are outside. This behavior can be accompanied by a rearrangement of the whole body without changing the general position of the behavior (Fig. 1B).

#### Rest Outside the Den

For this behavior, the octopus is outside the den without presenting movement activity, with the arms relaxed and extended around the body. This behavior can occur either at the bottom or on the walls of the tank and can be accompanied by a rearrangement of the whole body without changing the general position of the behavior (Fig. 1C).

#### Total Rest

The octopus is completely within the den with no visible activity and attached to one of the surfaces of its den (Fig. 1D).

### Locomotion behaviors

Behaviors were related to activity, characterized by different forms of movement of the octopus throughout the environment (Fig. 2).

**Figure 2.**
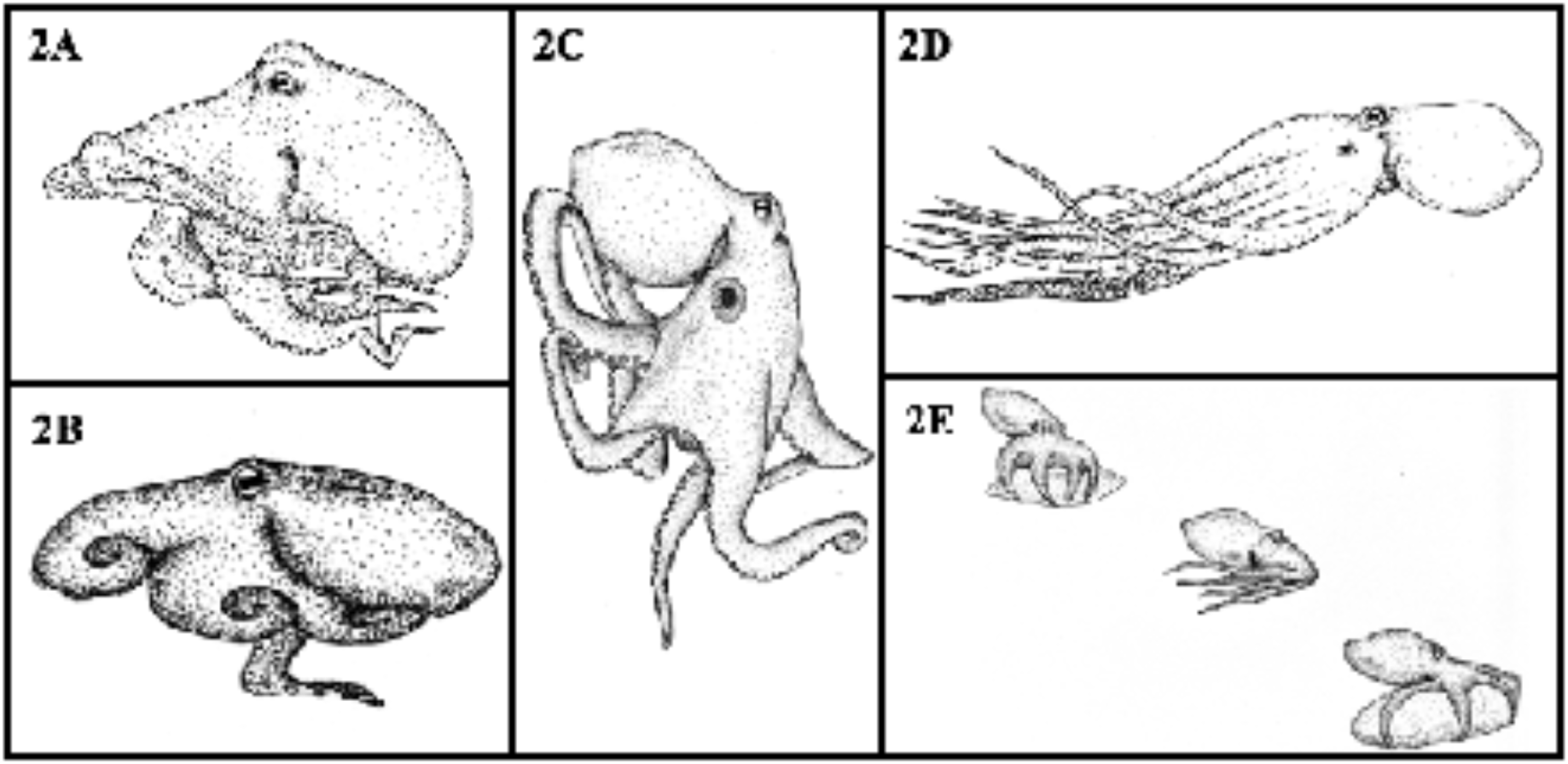
Locomotion behaviors of *O. maya*. A) Crawling: Type of locomotion in the bottom of the tank through the movement of the eight arms together. B) Hovering: The octopus moves in the bottom of the tank with only the fourth and fifth arms. C) Climbing: Locomotion along the walls of the tank in an ascending, descending, and horizontal way. D) Jet propulsion: The octopus makes use of a jet of water to move from one place to another. E) Jump on shells: Locomotion through jumping on shells and stones.

#### Crawling

Type of locomotion at the bottom of the tank through the movement of the eight arms together. This behavior occurs when the octopus leaves the den and explores the environment during periods of activity (Fig 2A).

#### Hovering

In this kind of locomotion, the octopus moves in the bottom of the tank with only the fourth and fifth arms, while the rest of the arms are partially coiled towards the back of the body and do not touch the bottom. This locomotion occurs when the distance to be traveled is short (Fig 2B).

#### Climbing

In this one, the octopus moves along the walls of the tank in an ascending, descending, and horizontal way, extending all arms to move. This behavior occurs when the octopus leaves the den and explores the environment during periods of activity (Fig. 2C).

#### Jet Propulsion

The octopus makes use of a jet of water through the siphon to move from one place to another more quickly. In this behavior, the position of the body of the octopus is different, since the mantle goes forward, and the arms are at the back. In general, this behavior occurs when the octopus needs to escape or approach another organism (Fig. 2D).

#### Jump on shells

The octopus moves in the tank by jumping, using the shells or stones that are close to each other as a surface and positioning itself on top of them (Fig. 2E).

### Feeding behaviors

These behaviors refer to the procedure of getting, ingesting, and eliminating food (Fig 3).

**Figure 3.**
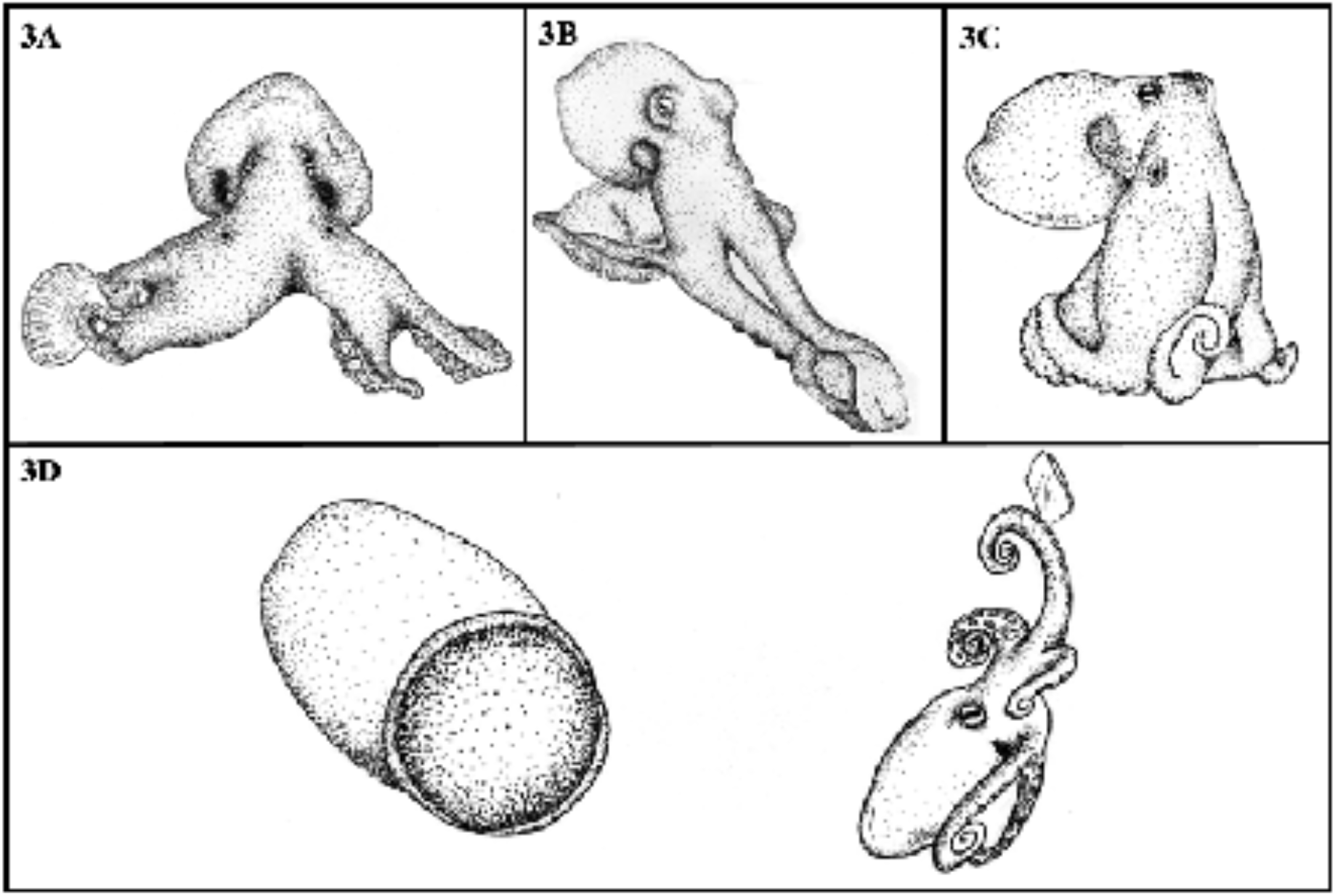
Feeding behaviors. A) Hunt: The octopus moves towards the prey with at least two arms and a propulsive movement. B) Reach: The octopus took the food extending and coiling the frontal arms without leaving its den. C) Feed: The interbrachial membrane covers the food, causing an elevation of the entire body. D) Waste Disposal: The octopus carries uneaten food away from its den

#### Hunt

When the octopus is fed, it moves towards the prey with at least two arms and a propulsive movement, reaching the food, wrapping it with the interbrachial membrane, and taking it with the beak. This behavior is most often seen when the octopus is fed with living food, however, it can also occur with frozen food (Fig. 3A).

#### Reach

This behavior occurs specifically when something is approaching to the octopus (for example frozen food). In the feeding procedure with frozen food, a stick is used for presenting the food which is taken by the octopus extending and coiling the frontal arms without leaving its den, allowing it to detach the food from the stick and carry it to the beak (Fig. 3B).

#### Feed

Once the octopus takes the food by displaying some of the previous behaviors (hunt; reach), it acquires a feeding posture characterized by an accelerated contraction and relaxation of the mantle. Besides, in this posture the interbrachial membrane and the base of the eight arms are covering the food, causing an elevation of the entire body but the distal part of the arms, which are relaxed close to the body. The octopus stops this behavior until the end of the feeding (Fig. 3C).

#### Waste Disposal

When the feeding behavior has been done, **t**he octopus carries uneaten food away from its den. The behavior can occur whether by wrapping the food with the interbrachial membrane or transporting away from the den by one of the aforementioned locomotion behavior. Or, using the arms to move away the remains of food without leaving the den, and even with the help of a jet of water expelled from the siphon moves it away even more (Fig. 3D).

### Home Maintenance

Behaviors related to the modification and cleaning of the den (Fig. 4).

**Figure 5.**
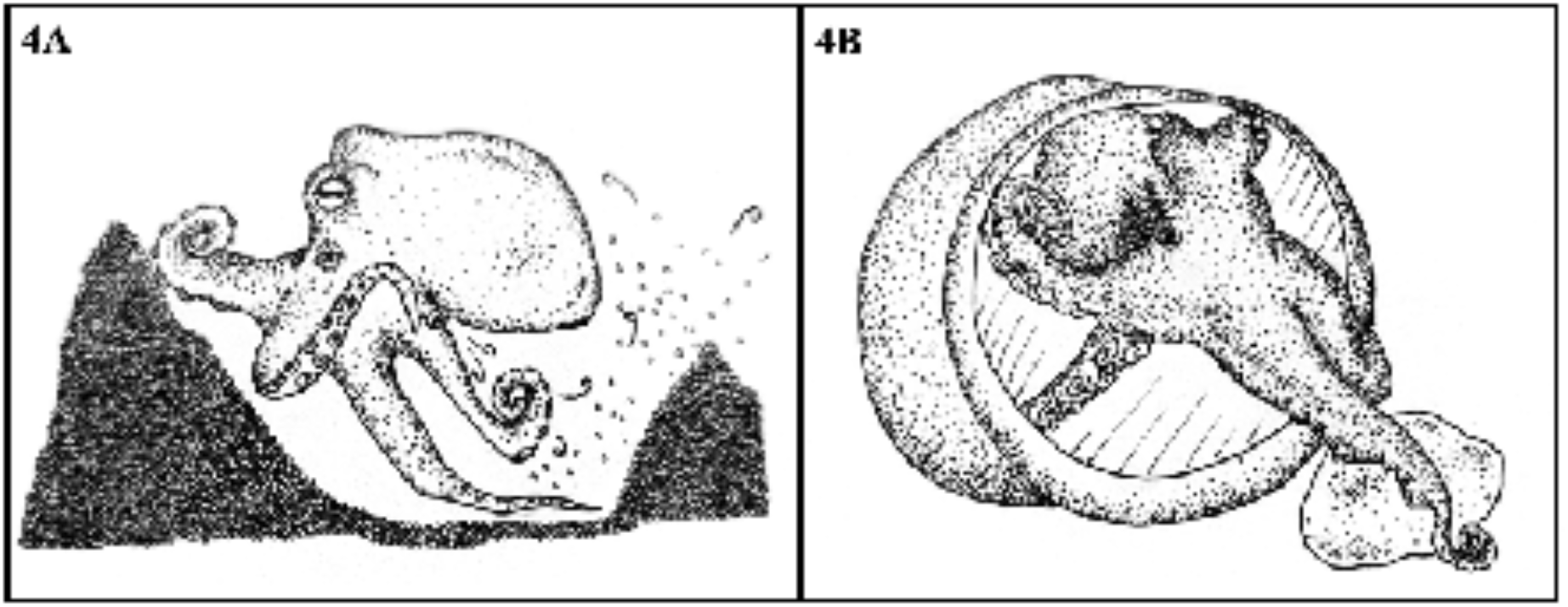
Home maintenance behaviors. A) Excavation: The octopus uses the siphon to remove sand under its body. B) Manipulate: The octopus uses one of its arms to hold various objects.

#### Excavation

In this behavior, we observe that the octopus uses the siphon to remove sand from under its body. It also uses the back arms to move the sand and stones towards the frontal arms to remove them from the area to make a hole in the bottom of the tank (Fig. 4A).

#### Manipulate

The octopus uses one of its arms to hold various objects through the suckers (e.g. shells or stones), either to bring them closer, move them away, or rearrange them inside or at the entrance of the den (Fig 4B).

### Protective behaviors

Behaviors related to the avoidance of aversive stimuli (Fig. 5). These behaviors were only seen during the full cleaning procedure of the tanks.

**Figure 5.**
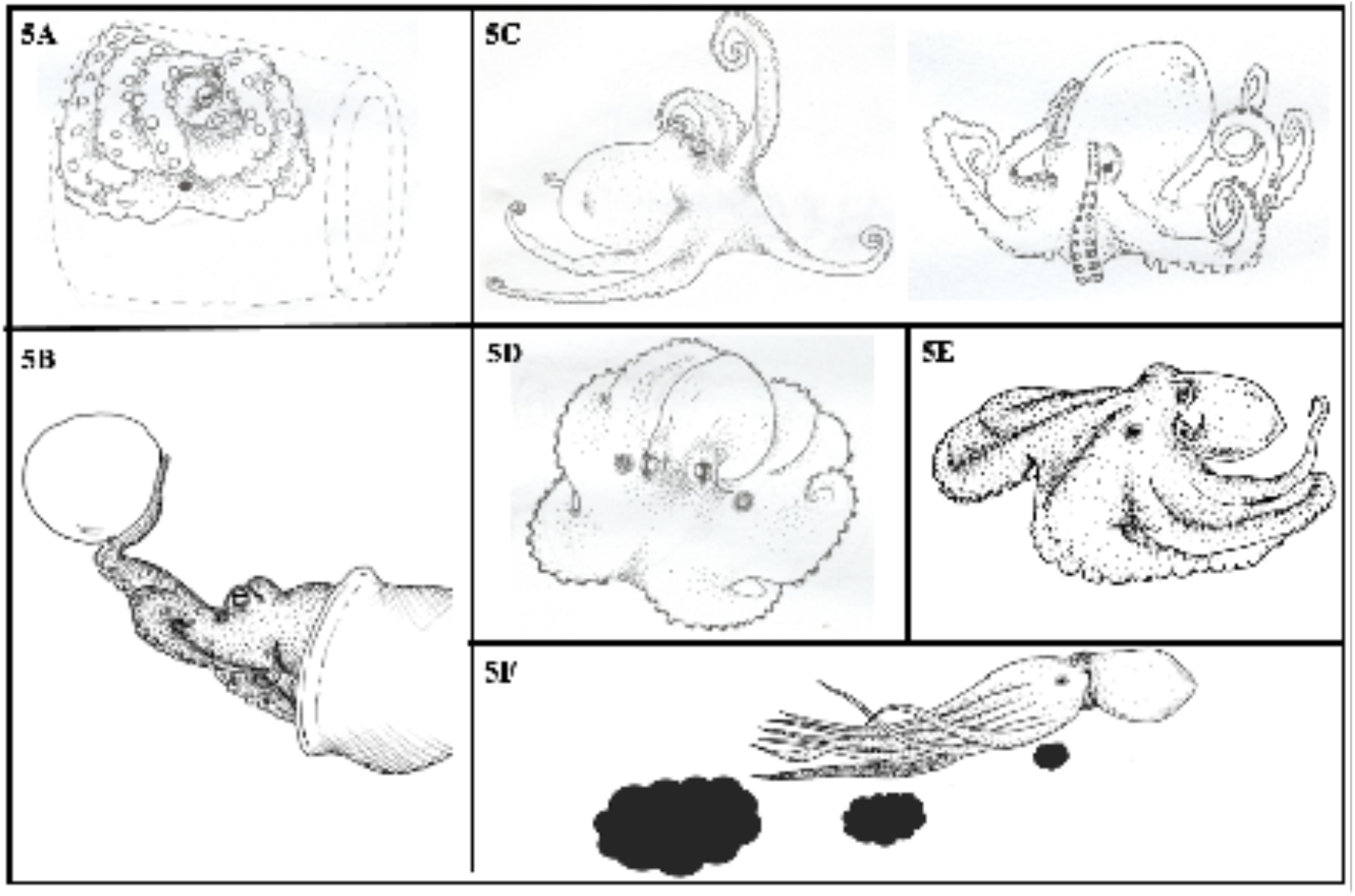
Protective behaviors. A) Isolation: The octopus is completely inside its den and exhibits an anchored posture. B) Push: The octopus uses one of its arms to move an object away from its body. C) Protection: The octopus uses its arms to protect the head or/and mantle from an aversive stimulus. D) Hand fan: The octopus’ interbrachial membrane is extended on the surface of the tank. E) Flatten: The entire body of the octopus is closely attached to a surface and spread out, without burrowing into the substrate. F) Inking: The octopus releases a jet of ink through the siphon.

#### Isolation

The octopus is completely inside its den and exhibits an anchored posture, characterized by the adherence of the eight arms to the surface of the den, completely surrounding the mantle and head. Only the beak and one eye are left exposed. This behavior is triggered by an aversive stimulus but is also related to senescence (Fig. 5A).

#### Push

The octopus uses one of its arms to move an object away from its body. This was observed mainly during feeding when the octopus repeatedly rejected food by moving away from the feeding instrument (Fig. 5B).

#### Protection

The octopus uses its arms to protect the head or/and mantle from an aversive stimulus, exposing the front part of the body, partially or totally, by bending the front arms or by curving all of them (Fig. 5C).

#### Hand fan

The octopus’ interbrachial membrane is extended on the surface of the tank, the base of the arms is slightly arched and the ends are curved in the same direction. The head is slightly elevated and stands out with respect to the rest of the body. This behavior is triggered by an aversive stimulus and sometimes is present with locomotion and rest behaviors (Fig. 5D).

#### Flatten

The entire body of the octopus is closely attached to a surface and spread out, without burrowing into the substrate. The arms can be extended on the surface or retracted close to the body. This behavior is triggered by an aversive stimulus and sometimes is present with locomotion and rest behaviors (Fig. 5E).

#### Inking

The octopus releases a jet of ink through the siphon, which is triggered by an aversive stimulus (e.g., a predator) and occurs in conjunction with the jet propulsion behavior, allowing a rapid escape movement (Fig. 5F).

### Other behaviors

Other behaviors do not fall into the above categories (Fig. 6).

**Figure 6.**
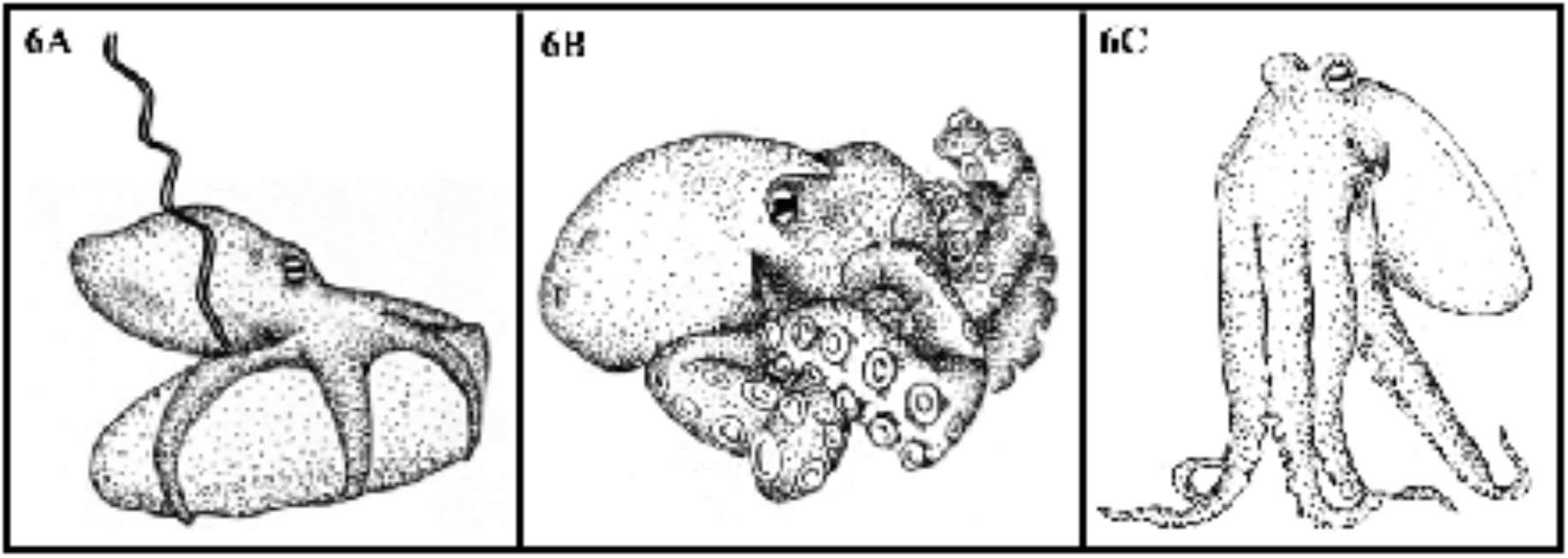
Other behaviors. A) Defecation: Expulsion of organic waste through the siphon. B) Grooming: The octopus coils all its arms towards the mantle and head, rubbing them in circular movements. C) Rearing: The octopus put together the eight arms, extends them, and rests on their tip.

#### Defecation

Expulsion of organic waste through the siphon. This behavior is very quick and does not usually last more than a couple of seconds (Fig. 6A).

#### Grooming

The octopus coils all its arms towards the mantle and head, rubbing them in circular movements. It’s a behavior related to the cleaning of the octopus itself, such as the elimination of residues, sand, and the layer of chitin that covers the suckers. (Fig. 6B).

#### Rearing

The octopus put together the eight arms, extends them, and rests on their tip, increasing its height and separating the rest of its body from the surface. This behavior can be present at the same time with locomotion and rest behaviors (Fig. 6C).

## OBSERVED BEHAVIORS OF EACH OCTOPUS FOR 24 HOURS

Using the categorization of behaviors previously described for *O. maya*, the appearance of each of all these was recorded for 24 hours continuous in four different specimens (Annex. Table 2).

The presence of all behavioral categories during the 24 recorded hours was observed on all specimens. However, the protective behavior category was not observed during the continuous recording of the day, this result looks promising due to this kind of behavior are related to stress and discomfort, meaning that our laboratory conditions seem to be right for the maintenance of the octopuses. Additionally, all specimens spent the highest percentage of time in non-activity-related behaviors (rest within the den, rest outside the den, and total rest), while activity-related behaviors showed a lower percentage in comparison (feeding, locomotion, den maintenance, and other behaviors).

## DISTRIBUTION OF BEHAVIORS AND PERIODS OF ACTIVITY (DAY AND NIGHT)

In order to identify variations in the distribution of behaviors during the day with respect to the night, the total times of all behaviors within the 24 hours were grouped into 12 hours corresponding to the day (08:00-19:00 hrs.) and 12 hours corresponding to the night (20:00-07:00 hrs.) based on the standard conditions of light and darkness from housing inside the laboratory. During the day, the presence of resting behaviors was demonstrated in 80.44%, locomotion behaviors in 10.18%, feeding behaviors in 8.85%, den maintenance behaviors in 0.24%, and other behaviors in 0.30% (Figure 7A). On the other hand, at night the presence of resting behaviors was observed in 94.85%, locomotion behaviors in 2.12%, feeding behaviors in 2.58%, den maintenance behaviors in 0.17%, and other behaviors in 0.28% (Figure 7B).

**Figure 7.**
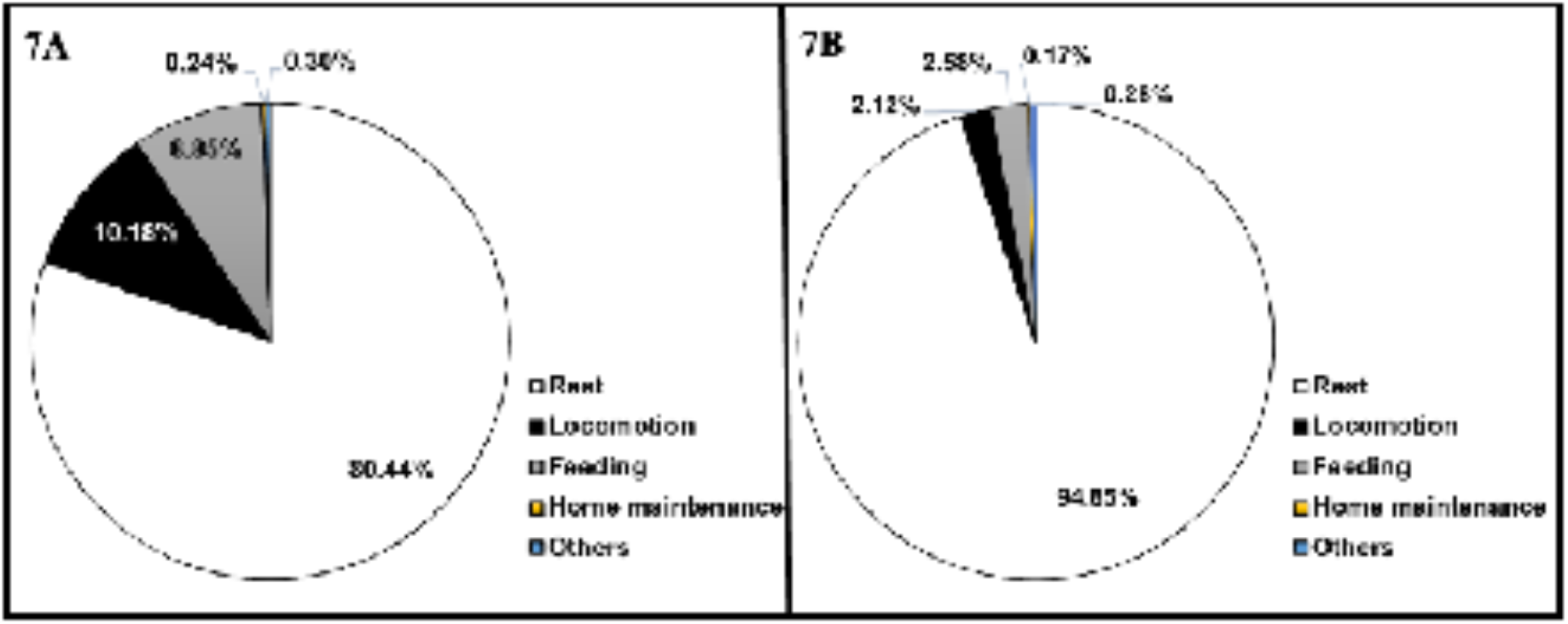
Distribution of behaviors along the day. A) Distribution of the behaviors that all the specimens present during the light hours. B) Distribution of the behaviors that all the specimens present during the dark hours. The data is presented as a percentage of the mean.

Furthermore, in order to obtain the activity periods of *O. maya*, the diurnal and nocturnal activity of the previously grouped 24 hours data was compared. The non-parametric Wilcoxon signed-rank test was made considering the data of all individuals, obtaining statistically significant differences between day and night (0.000602, p < 0.05) (Figure 8). This indicates that periods of activity during the 24 hours are differentiated. Taking together these results, with the ones of distribution of behaviors, demonstrate a higher activity present during the day compared to the activity during the night.

**Figure 8.**
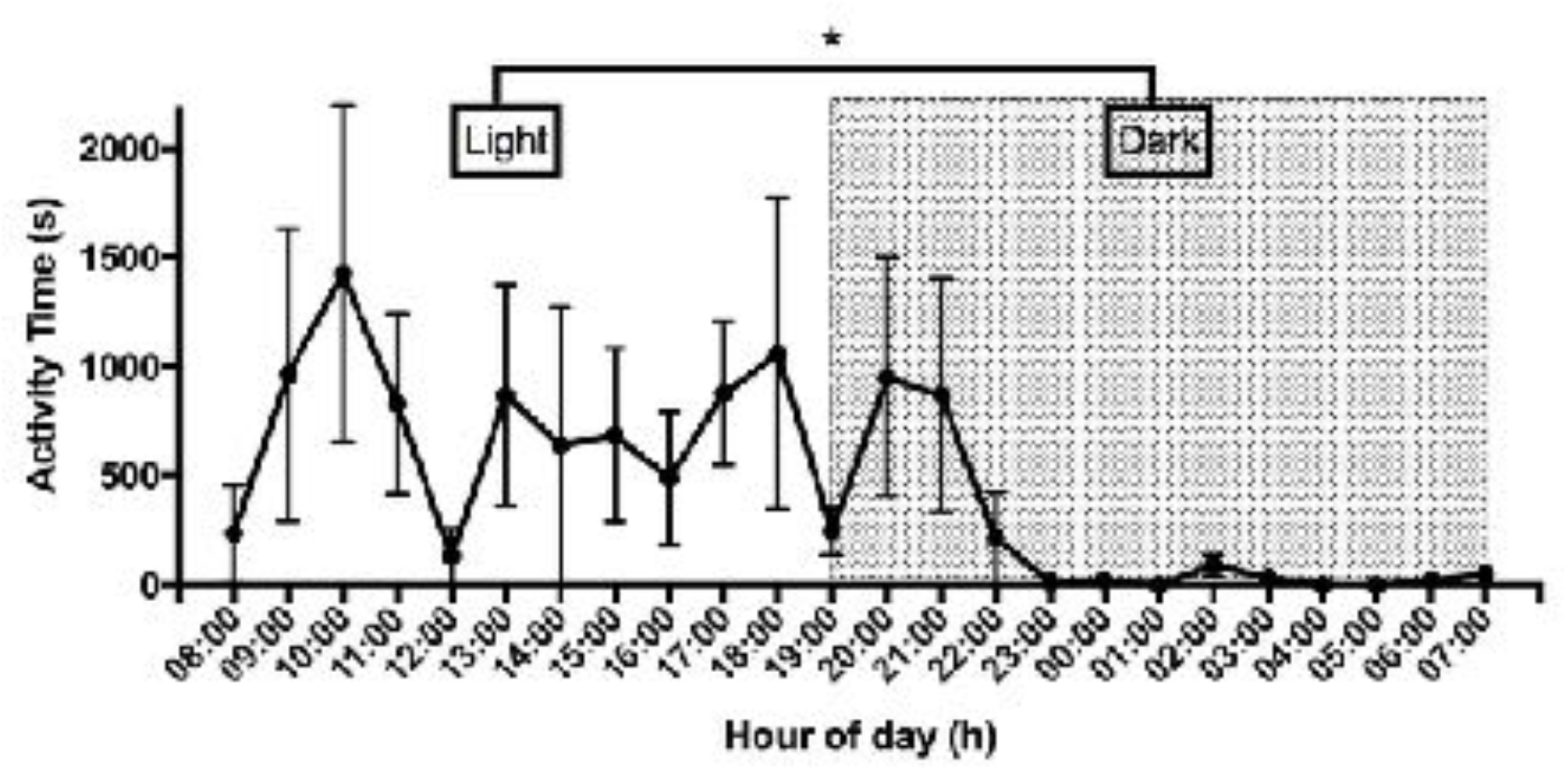
Activity time of the four specimens per hour (24 hours), during the light hours (08:00-19:00), and dark hours (20:00-07:00). Data is presented as mean and standard error. Significance was marked with * (P < 0.05)

## FREQUENCIES OF ACTIVITY BEHAVIORS

In addition to time, the frequency was also recorded for the four specimens during the 24 hours for activity-related behaviors (grooming, excavation, defecation, manipulation, crawling, hovering, jet propulsion, climbing, and jumping on shells), to obtain behaviors that occur most frequently in *O. maya*. Non-parametric Friedman`s ANOVA test proved that the means of activity behaviors were different (p < 0.05). Subsequently, Dunn’s multiple comparison test was made to determine the differences between each of the behaviors. Grooming behavior was statistically different against hovering (p < 0.01), excavation (p < 0.05) and jump on shells (p < 0.01); Manipulate behavior was statistically different against hovering (p < 0.05) and jump on shells (p < 0.05); crawling was statistically different against jump on shells (p < 0.05) and climbing was statistically different against jump on shells (p < 0.05) (Table 3). The behaviors with higher observed frequencies were grooming (mean = 22.75), crawling (mean = 26), and climbing (mean = 36) behaviors (Figure 9).

**Figure 9.**
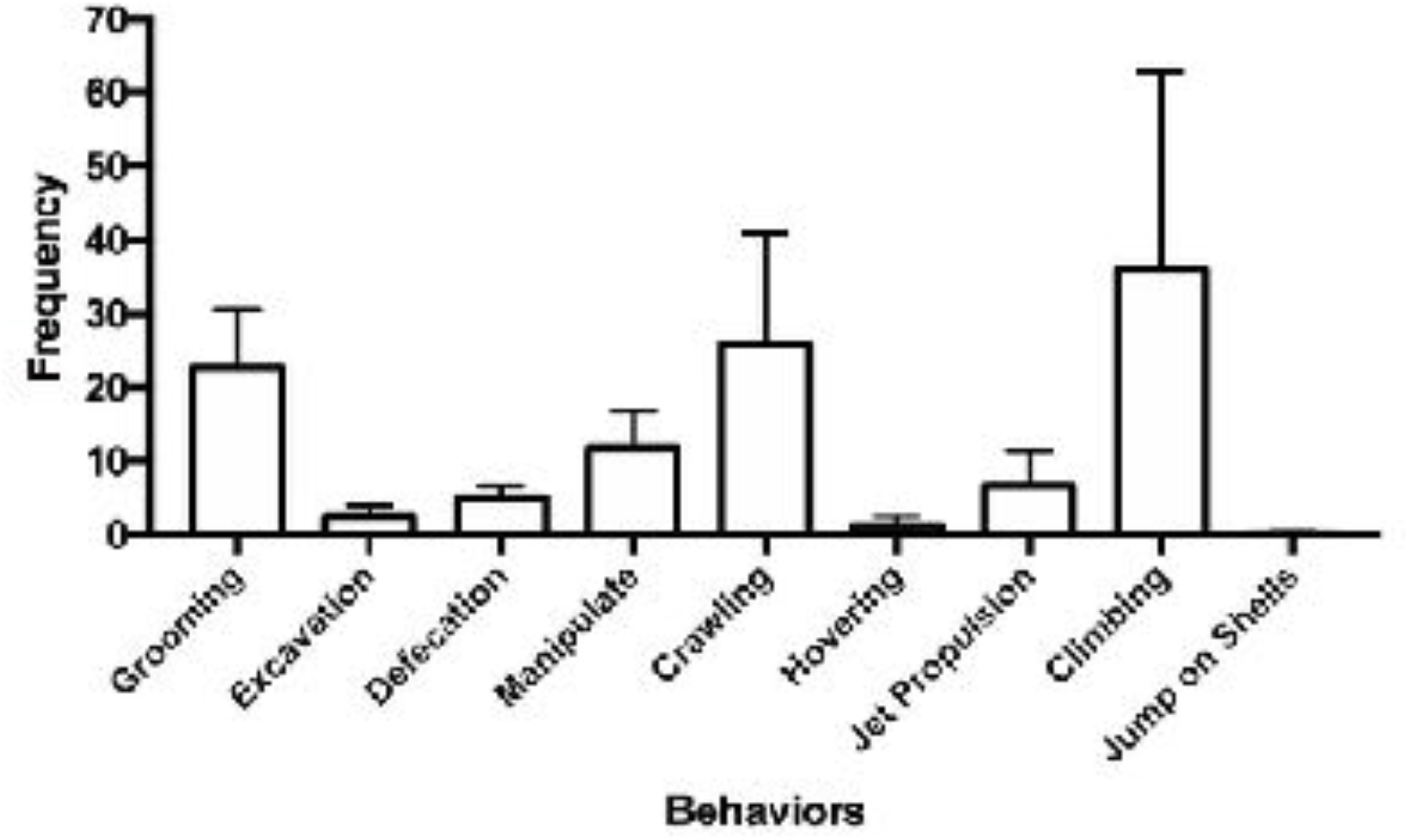
Mean frequency of behaviors related to activity during the 24 hours. Grooming, crawling, and climbing showed higher frequencies.

## DISCUSSION

### Behaviors of *O. maya*

Often, literature on octopus behavior mainly focuses on anecdotal reports of different species, and exhaustive description is quite rare (Borrelli & Fiorito, 2008; Marini et al., 2017). This poses an important problem since the evidence obtained on this model hasn’t had enough experimental control and results vary a lot. In other species of cephalopods such as cuttlefish, displayed behaviors have been extensively described, mainly in *Sepia officinalis* (Hanlon & Messenger, 1988) and *Sepia pharaonis* (Nakajima & Ikeda, 2017), being a benchmark for understanding the behavioral ecology of the genus Sepia and allowing quantifiable behavioral studies in these organisms (Nakajima & Ikeda, 2017). Just as the ethograms have helped other research models, this detailed ethogram intends to be used as a tool for behavioral quantification in other investigations and for them to be replicable in any laboratory without the need for special equipment or techniques.

There are few ethograms of different octopus species, which makes difficult the comparison between behavioral repertoires. However, in general, it has been reported that some postures and behaviors displayed by *O. maya* are also exhibited by other members of the Octopodidae family (Mather & Alupay, 2016; Huffard, 2007).

Regarding behaviors of the rest category, the rest within the den described in *O. maya* is very similar to that described in young octopuses of *O. vulgaris* in their wild environment, in which they remain a large percentage of daylight hours inside their den or some other safe place (Mather, 1988). In addition, *O. vulgaris* also presents inactivity periods throughout the day under laboratory conditions, characterized by presenting sporadic movements of the arms or body, but always remaining in the den (Meisel et al., 2011), similar to here described in *O. maya* during rest within and outside the den. Huffard (2007) also reports resting behaviors in *A. aculeatus* in wild environments, in which the octopus remains at the entrance of the den with its eyes raised and with the front part of the body exposed, similar to that seen in *O. maya*. Rest outside the den has also been described in this octopus, but there aren’t any reports about the presence of this behavior in other species of octopus. A possible explanation for this characteristic in *O. maya* could be the absence of predators in the laboratory conditions, since there is no danger of encountering an aversive stimulus in the environment, allowing the octopus to remain static outside its den for long periods of time, something that in the wild is unlikely to happen because the probability of survival would decrease. However, some reports indicate that *O. vulgaris* in the wild presents a feeding behavior outside the den, which means that it remains immobile for a period of time away from the den until feeding is complete (Mather, 1991a; Mather, 1991b), indicating that there are occasions when octopuses may present immobility outside the den. On the other hand, the total resting behavior observed in *O. maya*, in which the octopuses are completely introduced into their den and present inactivity for a long period of time, has been previously observed in *O. vulgaris* in the wild and in laboratory conditions and has been cataloged as a sleep-like behavior (Mather, 1988; Meisel et al., 2011). However, due to the limitations of the study, it was not possible to observe in detail this behavior while they remained inside the den, hence further experiments need to be done to corroborate and characterize the presence of sleep states in this species and its relationship with total resting behavior.

Regarding locomotion behaviors, five different types of locomotion were observed in *O. maya*. The first is crawling and it’s one of the most common behaviors present in all species of benthic octopus (Mather, 1998; Mather & Alupay, 2016), however, this behavior may present slight differences depending on the species. For example, in the case of *A. aculeatus* (Huffard, 2006) and *A. marginatus* with the description of so-called bipedal locomotion (Huffard et al., 2005). These differences in locomotion behaviors between species may explain the presence of hovering, climbing, and the jump shells behaviors found in *O. maya*, behaviors not previously described in other octopuses and it could be specified in this octopus species. On the other hand, the propulsion behavior has been described in the species *A. aculeatus* (Huffard, 2006), *O. vulgaris* and *O. insularis* (Mather & Alupay, 2016) in a very similar way, keeping the arms united to each other to move quickly from one place to another, although this same behavior has also been observed with a more pronounced separation between the arms as in *E. dofleini* (Mather & Alupay, 2016). Hence, the emphasis on a detailed study of octopus’s behavior gains relevance because all known species of octopus live in a variety of marine habitats, conditions that shape the complexity of the central nervous system and visual pathways (Chung et al., 2022) and also could shape these differences in behavioral performance between octopus’ species.

The process of feeding behavior reported in *O. maya* is very similar to that described in *O. vulgaris*, both species of octopus have a reaching behavior with their arms towards the food (called “poke” in *O. vulgaris*), and both present the same ingestion procedure by transporting the food through the arms towards the interbrachial membrane and maintaining it in the mouth until the feeding is done, finally, they also present behavior of elimination of remaining food through a jet of water or transported away from the den by the octopus itself (Mather, 1991a). In the same way, behaviors of the home maintenance category are also behaviors widely described in *O. vulgaris* (Mather, 1988; Mather, 1994) described as a modification of the den by removing sand from the substrate or placing rocks and objects in front of this, in a very similar way to that observed in *O. maya*. In this sense, excavation and manipulation behaviors used to change the distance of objects and to dig in the sand reported here, have also been previously described in *O. vulgaris* (Mather, 1994; Mather, 1998; Mather & Alupay, 2016). Even this great motility and variety of movement of the eight arms have already been characterized previously (Mather 1998; Mather & Alupay, 2016). Finally, behaviors such as grooming and defecation have also been widely reported in *O. vulgaris* (Packard & Sanders, 1971; Mather, 1988, Mather & Alupay, 2016) and in *A. aculeatus* (Huffard, 2007), and the behavior of rearing has also been described in *O. vulgaris, O. cyanea, Enteroctopus dofleini* (Mather & Alupay, 2016) and *A. aculeatus* (Huffard, 2007) and the performance of these behaviors are not usually very different from those observed in *O. maya*.

On the other hand, it was only possible to observe protective behaviors during and after the full cleaning procedure of the experimental tanks, in which the octopus was taken out of their tanks and put in a smaller tank with water from the system, this for no more than 10 min while the cleaning procedure was made. Although the procedure was carefully performed and avoided the stress on animals, it has been reported that just handling and exposure to air for short periods of time is enough to generate a state of stress in octopuses (Malham et al., 2002). In this sense, it was only during this procedure that we observe protection behavior in *O. maya* characterized by a curl of one or all the arms, a body pattern previously reported in young (Packard & Sanders, 1971) and adult specimens of *O. vulgaris* (Mather, 1998). In both species, it was associated with a defensive response to aversive stimuli like the threat of predators and the presence of other octopuses. In the same way, these postures present a certain similarity to the “Flamboyant” posture, a stereotyped cryptic posture that is characterized by a helix-like twisting of the arms towards the mantle and head and the lifting of the body from the substrate, which has been widely described in young octopus’ species: *O. vulgaris, E. dofleini, A. aculeatus, O. chierchiae, O. gorgonus and O. joubini* (Huffard, 2007, Mather & Alupay, 2016; Mather, 1998; Packard & Sanders, 1971). However, the only difference between the “Flamboyant” posture and the one seen in *O. maya* is that the raised papillae seen in the aforementioned species are not present in *O. maya*. On the other hand, the inking behavior is one of the most common behaviors in members of the Octopodidae family, so no differences were found in the performance of these behaviors between *O. maya* and other octopus’ species (Mather & Alupay, 2016). The flat behavior has been described in *O. vulgaris, O. cyanea, Enteroctopus dofleini* (Mather & Alupay, 2016), and *A. aculeatus* (Huffard, 2007), and hand fan behavior has also been reported in *O. vulgaris, O. briareus* and *Thaumoctopus mimicus* (Packard & Sanders, 1971; Mather & Alupay, 2016; Norman & Hochberg, 2005). Lastly, for the other two protective behaviors, isolation and pushing behavior were also only observed after the cleaning procedure, when the specimens are returned back to their individual tanks, but these behaviors did not remain in time. Unfortunately, there are not so many studies where the stress response of octopuses is evaluated; therefore, it’s difficult to compare with other species in this behavioral category. Besides, it is important to mention that protective behaviors were only observed during the *ad libitum* sampling and not during the analysis of the long 24-hour videos, this shows that the conditions in our facilities in which octopuses were housed are adequate as in order to not present behaviors related to aversive stimuli or environmental stressors and the descriptions of these stress-related behaviors could help to start to understand the behavioral stress response in octopus and prove the effects of different stressors in neurobiology and behavior.

It’s worth mentioning that although postures and behaviors described above have already been observed in many other species of octopus, they may be unreported or ignored because the behavioral assessment was not the main goal of these investigations or because of the absence of controlled experimental observations (Borrelli & Fiorito, 2008; Marini et al., 2017). For these reasons, the realization of this ethogram was not only for the description of behaviors that conforms to the behavioral repertoire of *O. maya* but also to address the quantification of these. Cephalopods are excellent neuroscience models because of their anatomical features and nervous system organization, behavior must be equally important than this because of the wide flexibility of behavioral repertoire. Hence, this study aims to encourage behavioral evaluation in octopus research since it allows us to have a greater notion and control of the distribution and frequency of these organisms’ behaviors and, consequently, to be able to observe changes under different conditions and have a better brain-behavior neuroscience study.

### Distribution of resting behaviors and periods of activity (day and night)

*O. maya* spent a high percentage of their time throughout the day in behaviors related to resting, presenting them 80.44% of the time during the day and 94.85% during the night. This agrees with the previously reported data stating that in general octopuses are known to present very long periods of rest during the 24 hours of the day, confined mainly to the den (Mather et al., 1985; Mather, 1988). Such is the case of juvenile octopuses of *O. vulgaris* in the wild, which spend 70% of daylight hours inside a den (Mather, 1988). The same could be observed in laboratory conditions in this same species, where it is reported that adults have a mean activity below 50% throughout the day (Meisel et al., 2003; Meisel et al., 2006). More recent laboratory studies have also shown that this species presents an average duration of 16.7 hours of inactivity during the 24 hours of the day, characterized by always remaining inside its den (Meisel et al., 2011). Similar results have also been found in other species, like *O. macropus* which presents an average activity below 50% during the high activity periods (Meisel et al., 2006), *O. insularis* presents 93.58% of inactivity-related behaviors throughout the day (Medeiros et al., 2021), in *O. laqueus* a global activity duration of 7 to 14 minutes has been reported during the day and 298 to 339 minutes during the night in 24 hours recording, and in *A. aculeatus* a global activity duration of 49 to 99 minutes during the day and 138 to 185 minutes during the night (Ikeda & Yanagisawa, 2018). The information above indicates that, like many other species of octopus, *O. maya* also presents prolonged periods of inactivity throughout the 24 hours of the day, something related to an efficient predator ability in these animals (Mather, 1988) and a good sign of acclimatization to the environment, even in a laboratory condition.

In regard to activity periods, it could be observed that *O. maya* exhibits greater activity during the day compared to the night in laboratory conditions, showing the presence of activity-related behaviors mostly during light hours (behaviors of locomotion, home maintenance, feeding, and others), distributed between 08:00 and 17:00 hrs. The activity periods in octopus species are still highly unclear, mainly because they differ depending on the species being studied, moreover, the variables to which octopus synchronize their activity is not very clear yet (Mather, 1988). There are even some reports in which different periods of activity occur in the same octopus’ species (Meisel et al., 2011). Such is the case of *O. vulgaris*, which has been reported with nocturnal activity (Kayes, 1974) but also reports with a diurnal activity (Mather, 1988), and in juvenile organisms, a twilight activity has been reported (Mather, 1991a). A possible explanation for this could be the fact that marine organisms’ biological rhythms are influenced by environmental variables such as temperature, light, or tides (Ikeda & Yanagisawa, 2018); variables that fluctuate between oceanic regions, and likely cause the contrast in the found evidence.

Results in *O. maya* are similar to those seen in *O. cyanea*, characterized by starting activity at 07:00 and reaching a peak between 13:00 and 18:00 (Houck, 1982). Data is also consisting with the results obtained in some studies carried out on *O. vulgaris* in the wild, where it was reported that it has greater activity during the day and occupancy of the den at night (Mather, 1988), moreover, this kind of diurnal activity presented in this species has also been confirmed in a laboratory environment (Meisel et al., 2006). On the other hand, results obtained in *O. maya* contrast with those observed in *E. cirrhosa* which are described as nocturnal species with greater activity during periods of reduced illumination (Cobb et al., 1995), also with *O. macropus* characterized by greater activity at night, in both wild and laboratory environments (Meisel et al., 2006) and finally contrast data of *O. dofleini* where it presents a slightly higher activity at night with activity peaks between 21:00 and 02:00 (Mather et al., 1985). A possible explanation for this discrepancy between species may be that *O. maya* is an intertidal species that move in shallow waters, while *E. cirrhosa, O. macropus*, and *O. dofleini* are octopus’ species that inhabit deep water, in which light and environment conditions are different (Cobb et al., 1995). Similarly, these factors would also explain the concordance with what is seen in the species *O. vulgaris* and *O. cyanea* because both are shallow water species like *O. maya* and maybe the same environmental variables controlled the activity periods. However, more experiments are needed to clarify these differences in the activity periods between octopus species to find the main exogenous environmental factors or an endogenous biological clock that synchronized their activity periods.

### Frequency of behaviors

The study of behavior through more controlled methods is something that has recently begun to be addressed, through the proposal of specific behaviors for research, like in the study of *O. vulgaris* and the use of attack behavior as a parameter to evaluate in behavioral analysis (Borrelli et al., 2020). However, as we see not all the behaviors are exhibited in the same way in other octopus’ species due to the differences given by environmental factors, so the behavior to choose as a behavioral parameter must always be adapted to the characteristics of the studied species. Knowing the behavioral repertoire of octopus species gives a list of behaviors to choose from according to the purpose of research and the quantification of these help to characterize the differences between species. These features give better control to choose a behavioral parameter in the analysis of octopus behavior. In this sense, this ethogram and the reporting of frequency distribution in *O. maya* throughout the day allows recognizing the behaviors that occur most frequently in this species, being climbing, crawling, and grooming behaviors with greater frequency of appearance. These data allow us to propose these behaviors as a behavioral parameter for the analysis of *O. maya* in tasks related to cognition and behavior given their high frequency of appearance compared to others. However, since there were few octopuses for use in this study, further studies are required to corroborate the information.

## CONCLUSIONS

According to the results obtained in this research, the octopus species *O. maya* bred in captivity and kept in laboratory conditions demonstrates a wide repertoire of behaviors, exhibiting at least twenty-three behaviors distributed in six different categories during the behavioral analysis performed. In addition, this species shows periods of differentiated activity during the day and at night, showing activity peaks, distribution, and frequencies of behaviors related to activity mainly during daylight hours compared to night.

The decision of using an octopus species as a model depends on the purpose and objectives of the investigation but the initial works should be carried out with species that are easily available (Boletzky & Hanlon 1983). *O. maya* presents these characteristics because captive breeding in México is being developed successfully since 2004 by UNAM. Also, the adapted-well laboratory feature of *O. maya* gives a great advantage in the use of this species in neuroscience research, because the maintenance of the octopus in this condition for a long period is easier, even from hatching because the embryos hatch as holobenthic juveniles, animals with the same gross anatomical features as adults (Hanlon & Forsythe, 1985; Moguel et al., 2010; Rosas et al., 2007; Vidal, 2014), and as we see here, also presenting behaviors comparable with studies in other octopus species, demonstrate that behavior is stable in laboratory conditions.

These results showed that *O. maya* presents behaviors comparable with other octopus species and supports that it is an organism that can be proposed as a research model. Also, knowing the behavioral repertoire of this species allows greater control in subsequent investigations related to behavior and provides a great tool to constantly monitor the health status of organisms and its use for research, being a precedent for future behavioral research in this species related to neuroscience and cognition.

## SUPPLEMENTARY FILES

**Supplementary 1.**
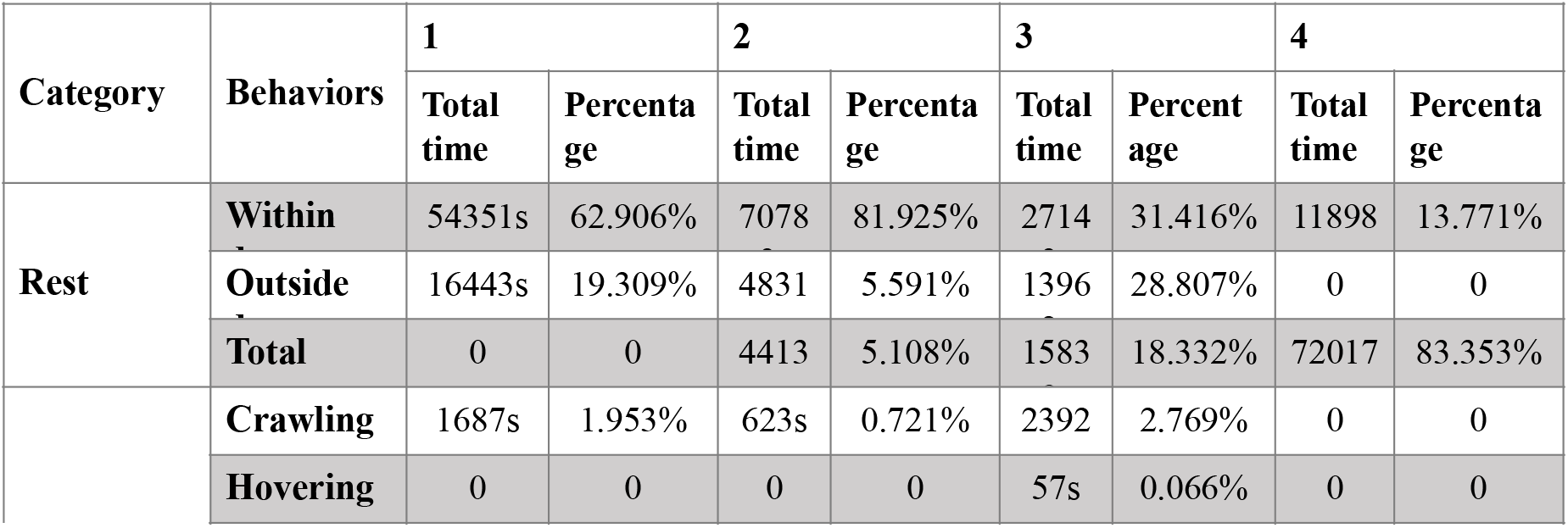
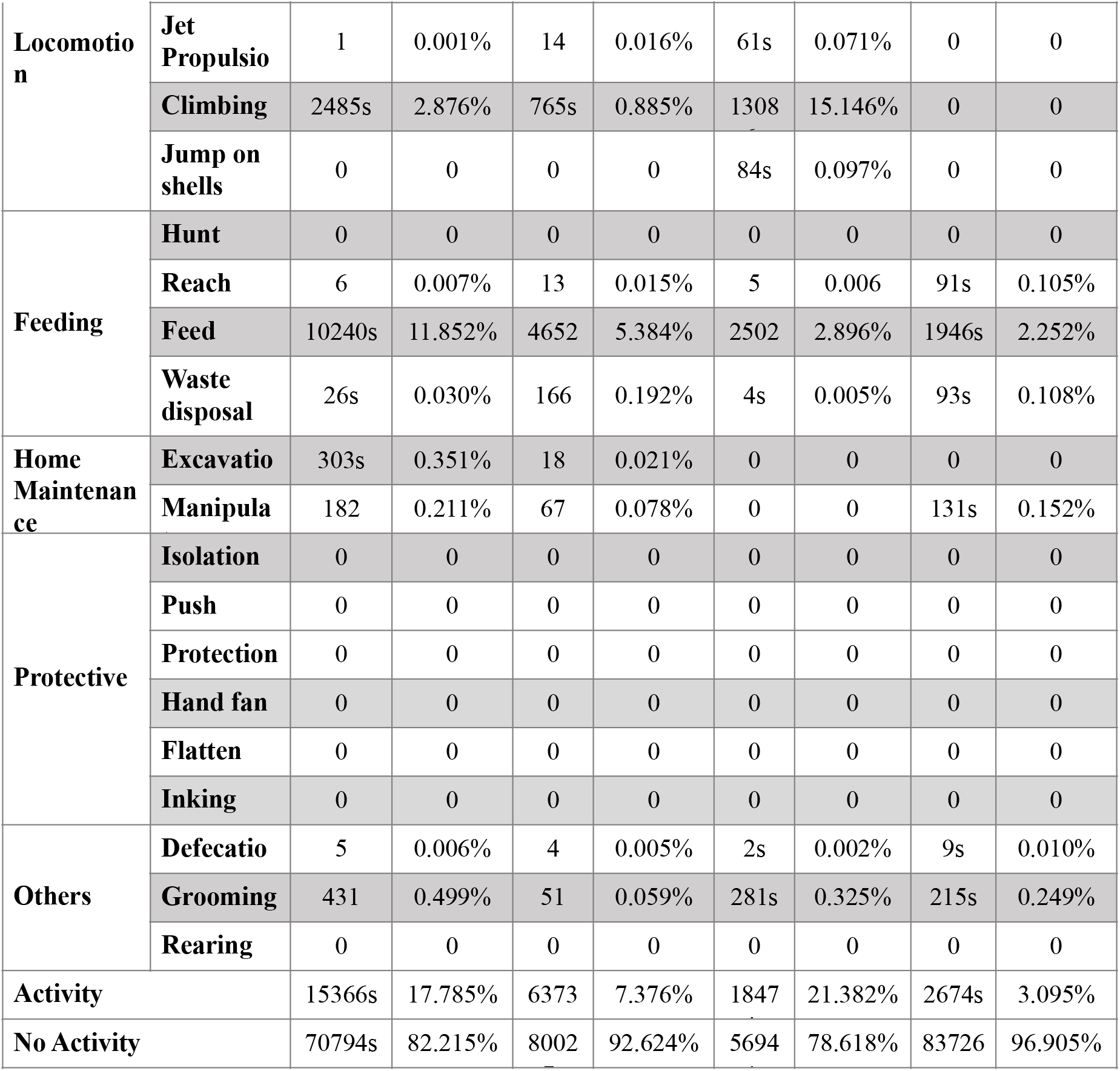
The total time and the percentage in 24 hours for each of the behaviors in four different specimens of *O. maya*. It’s worth mentioning that the protective behaviors present a 0% because these behaviors were only seen in the full cleaning procedure, in normal conditions, there is no aversive stimulus.

**Supplementary 2.** Results of uncorrected Dunn’s test, showing the significance and individual p value between each behavior. Significance (P<0.05).

| Uncorrected Dunn's test | Mean rank diff. | Significant? | Individual P Value |
| --- | --- | --- | --- |
| **Grooming vs. Excavation** | 16.25 | Yes | 0.0266 |
| **Grooming vs. Defecation** | 10.25 | No | 0.1619 |
| **Grooming vs. Manipulate** | 5.375 | No | 0.4633 |
| **Grooming vs. Crawling** | 5.5 | No | 0.4530 |
| **Grooming vs. Hovering** | 19.75 | Yes | 0.0070 |
| **Grooming vs. Jet Propulsion** | 12.5 | No | 0.0881 |
| **Grooming vs. Climbing** | 5.625 | No | 0.4428 |
| **Grooming vs. Jump on Shells** | 21.5 | Yes | 0.0033 |
| **Excavation vs. Defecation** | -6 | No | 0.4129 |
| **Excavation vs. Manipulate** | -10.88 | No | 0.1378 |
| **Excavation vs. Crawling** | -10.75 | No | 0.1424 |
| **Excavation vs. Hovering** | 3.5 | No | 0.6329 |
| **Excavation vs. Jet Propulsion** | -3.75 | No | 0.6089 |
| **Excavation vs. Climbing** | -10.63 | No | 0.1471 |
| **Excavation vs. Jump on Shells** | 5.25 | No | 0.4738 |
| **Defecation vs. Manipulate** | -4.875 | No | 0.5059 |
| **Defecation vs. Crawling** | -4.75 | No | 0.5169 |
| **Defecation vs. Hovering** | 9.5 | No | 0.1949 |
| **Defecation vs. Jet Propulsion** | 2.25 | No | 0.7588 |
| **Defecation vs. Climbing** | -4.625 | No | 0.5280 |
| **Defecation vs. Jump on Shells** | 11.25 | No | 0.1248 |
| **Manipulate vs. Crawling** | 0.125 | No | 0.9864 |
| **Manipulate vs. Hovering** | 14.38 | Yes | 0.0498 |
| **Manipulate vs. Jet Propulsion** | 7.125 | No | 0.3309 |
| **Manipulate vs. Climbing** | 0.25 | No | 0.9728 |
| **Manipulate vs. Jump on Shells** | 16.13 | Yes | 0.0278 |
| **Crawling vs. Hovering** | 14.25 | No | 0.0518 |
| **Crawling vs. Jet Propulsion** | 7 | No | 0.3395 |
| **Crawling vs. Climbing** | 0.125 | No | 0.9864 |
| **Crawling vs. Jump on Shells** | 16 | Yes | 0.0290 |
| **Hovering vs. Jet Propulsion** | -7.25 | No | 0.3225 |
| **Hovering vs. Climbing** | -14.13 | No | 0.0539 |
| **Hovering vs. Jump on Shells** | 1.75 | No | 0.8113 |
| **Jet Propulsion vs. Climbing** | -6.875 | No | 0.3482 |
| **Jet Propulsion vs. Jump on Shells** | 9 | No | 0.2194 |
| **Climbing vs. Jump on Shells** | 15.88 | Yes | 0.0303 |

